# Directed evolution of angiotensin-converting enzyme 2 (ACE2) peptidase activity profiles for therapeutic applications

**DOI:** 10.1101/2022.01.03.474817

**Authors:** Pete Heinzelman, Philip A. Romero

## Abstract

Angiotensin Converting-Enzyme 2 (ACE2) is currently being investigated for its ability to beneficially modulate the Angiotensin receptor (ATR) therapeutic axis to treat multiple human diseases, but its broad substrate scope and diverse physiological roles limit its potential as a therapeutic agent. In this work we engineer ACE2 variants with enhanced hydrolytic activity and specificity toward Angiotensin-II (Ang-II) that may lead to new therapeutic candidates with improved efficacy and reduced off-target side effects. We established a yeast display-based liquid chromatography screen that enabled use of directed evolution to discover ACE2 variants with improved Ang-II activity and specificity relative to the off-target peptide substrates Apelin-13 and Angiotensin-I (Ang-I). We screened ACE2 active site libraries to reveal three substitution-tolerant positions (M360, T371 and Y510) that can be mutated to enhance ACE2’s activity profile and followed up on these hits with focused double mutant libraries to further improve the enzyme. Relative to wildtype ACE2, our top variant (T371L/Y510Ile) displayed a sevenfold increase in Ang-II turnover number (*k*_*cat*_), a sixfold diminished catalytic efficiency (k_cat_/K_m_) on Apelin-13, and an overall decreased activity on other ACE2 substrates that were not directly assayed in directed evolution screen. At physiologically relevant substrate concentrations, T371L/Y510Ile hydrolyzes more Ang-II than wildtype ACE2 with concomitant Ang-II:Apelin-13 specificity improvements reaching 30-fold. Our efforts have delivered ATR axis-acting therapeutic candidates with relevance to both established and unexplored ACE2 therapeutic applications and provide a foundation for further ACE2 engineering efforts.

**Significance Statement:** The Angiotensin Converting Enzyme 2 (ACE2) carboxypeptidase is being clinically trialed for treatment of acute respiratory distress syndrome (ARDS) related to traumatic injuries and viral infections. We developed an enzyme engineering platform that enabled discovery of ACE2 variants possessing enhanced peptide hydrolysis activity and specificity profiles relevant to ARDS treatment. Beyond their potential utility as ARDS therapeutics, our improved ACE2 variants could be further developed for new unexplored ACE2 therapeutic contexts such as treating Alzheimer’s Disease.

## Introduction

Angiotensin-Converting Enzyme 2 (ACE2) is a zinc-dependent mono-carboxypeptidase found in both membrane bound and freely circulating forms that plays a central role in multiple peptide hormone signaling pathways in humans [1]. ACE2 cleaves the terminal phenylalanine residue from Angiotensin-II to produce Angiotensin (1-7), resulting in downstream vasodilation and increased salutary blood flow [2], suppressed inflammation [3] and attenuated pathological tissue remodeling after acute injury [2]. These beneficial physiological responses have motivated clinical researchers to pursue ACE2 as an agent for treating respiratory viral infections [4], acute respiratory distress syndrome [5], and diabetes [6].

While therapeutically administered ACE2 has great potential to modulate the Angiotensin-II (Ang-II) axis for treating human disease, this enzyme also acts on at least eight additional peptide hormones found in circulation and/or tissues [7,8]. Apelin peptides are major off-target substrates to consider due to their key role in the cardiovascular system, ACE2’s high catalytic activity towards multiple Apelin isoforms [7-9], and the fact that Apelin peptides are 1-2 orders of magnitude higher concentration than Ang-II in human blood and tissues [10-13]. ACE2-mediated inactivation of Apelins reduces their important cardioprotective effects [9] and could trigger myocardial dysfunction when administered therapeutically. ACE2’s broad substrate scope and distributed physiological roles greatly limit its potential to specifically cleave Ang-II for therapeutic applications.

In this work we engineer ACE2 variants with tailored substrate profiles to specifically cleave Ang-II while leaving off-target peptide signaling pathways such as those involving Apelins unaffected. We designed site-directed ACE2 libraries to target key substrate recognition pockets and applied liquid chromatography-based screening on mixtures of peptide substrates to mimic heterogeneous *in vivo* conditions and to identify variants possessing the desired substrate cleavage profiles. After two rounds of directed evolution, we identified ACE2 variants with substantially improved activity and specificity toward Ang-II over Apelin-13. Our leading variant (T371L/Y510Ile) displayed a sevenfold increase in its turnover number (k_cat_) toward Ang-II and also increased Ang-II:Apelin-13 specificity by up to 30-fold at physiologically relevant substrate concentrations. We also found our engineered ACE2s display high specificity for Ang-II relative to other peptide hormones that were not directly assayed in our library screening, suggesting a highly focused substrate scope that may allow independent modulation of the Ang-II therapeutic axis. The outcomes presented here will have an important impact on current and future clinical applications of ACE2 for the prevention and treatment of human disease.

## Results

### Yeast surface display-based directed evolution of ACE2 for enhanced activity and specificity

ACE2 acts on different peptide substrates to regulate multiple peptide hormone signaling pathways *in vivo* (Figure 1a). We aim to engineer ACE2 variants with focused substrate scopes to specifically modulate Ang-II levels. We developed a streamlined ACE2 screening platform and performed active site-targeted directed evolution to discover new ACE2 variants with enhanced activity and specificity toward Ang-II.

**Figure 1.**
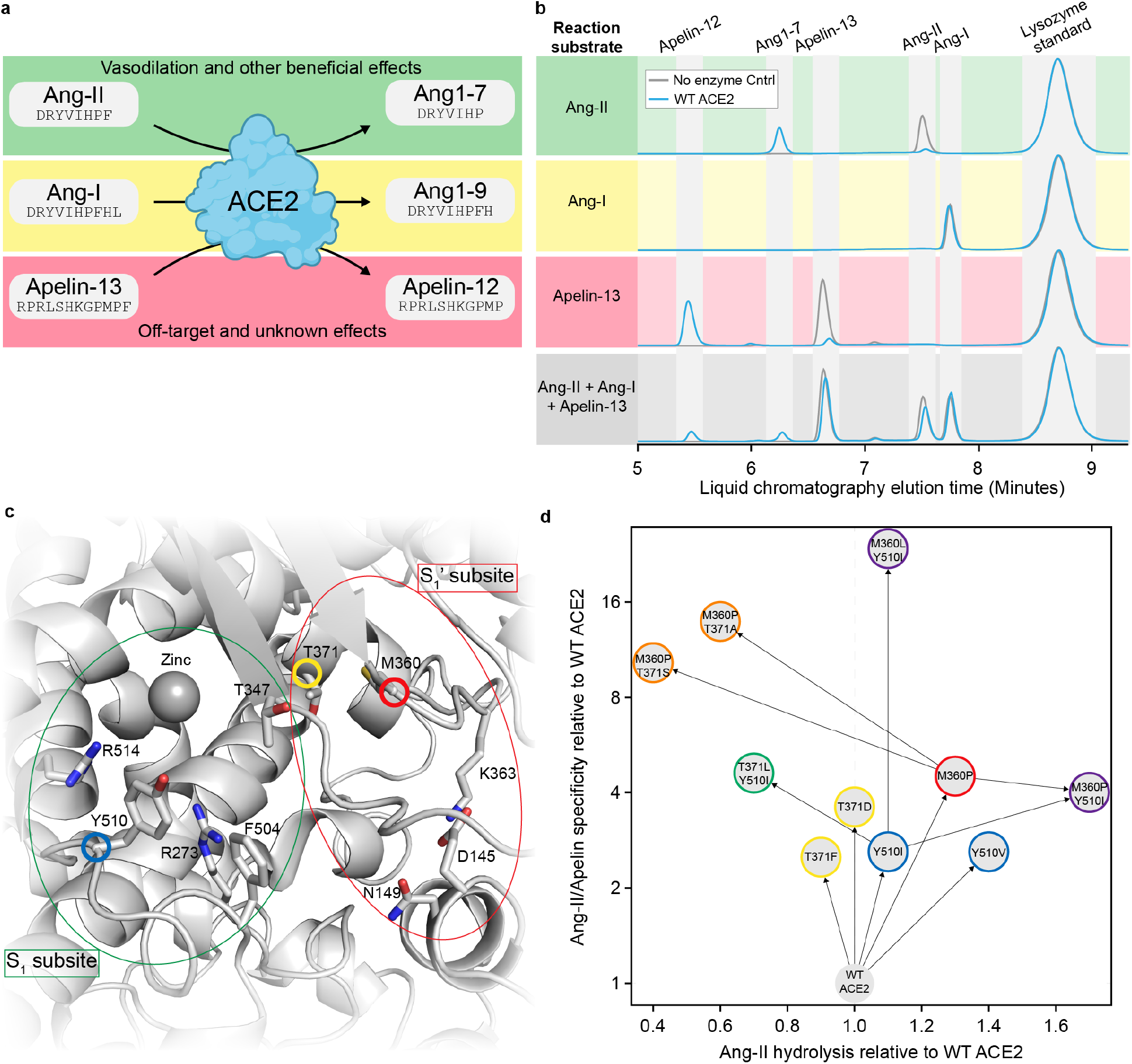
Directed evolution of ACE2 as a therapeutic agent. (a) Key ACE2 peptidase reactions relevant to the Ang-II/ATR therapeutic axis. Green background denotes hydrolysis reaction with beneficial *in vivo* effects. Red background denotes hydrolysis reaction with detrimental effects. Yellow background indicates unknown and potential for detrimental and/or beneficial, i.e., reduction in amount of Ang-I hydrolytic conversion to Ang-II by the Angiotensin Converting Enzyme (ACE) dipeptidase, effects. (b) LC chromatograms showing peptide hydrolysis by ACE2-displaying yeast. Our ACE2 system shows high peptidase activity on Ang-II and Apelin-13 substrates, and undetectable hydrolysis of Ang-I. We used horseradish peroxidase-displaying yeast as a negative control. (c) The ACE2 substrate binding pocket is composed of two subsites: S_1_ that encases the substrate residue N-terminal to the cleavage site and S_1_’ that accommodates the final C-terminal residue. The labeled residues correspond to positions for which we constructed and screened yeast-displayed NNB mutant libraries. The three circled residues indicate amino acid positions where mutations enhanced activity and/or specificity. (d) Overview of the directed evolution process. The Ang-II hydrolysis and specificity values were determined using mixed substrate assays from the YSD library screen. Specificity is defined as the ratio of moles Ang-II hydrolysis product to moles Apelin-13 hydrolysis product for the variant ACE2 divided by that same ratio for wildtype ACE2. The numbers reflect single or duplicate measurement from the YSD screen.

Our ACE2 screening platform leverages yeast surface display (YSD) [14,15] to express active ACE2 enzymes on the yeast surface (Supplementary Figures 1 and 2), allowing subsequent enzyme purification by simple centrifugation and resuspension into assay buffer. We then add individual peptides or mixtures of peptides to the ACE2-expressing yeast and monitor hydrolysis reactions via high performance liquid chromatography (LC). The combination of yeast display and LC-based screening enables rapid testing of ACE2 variants’ substrate hydrolysis profiles.

We tested our yeast display-LC screening platform on Ang-II, Angiotensin-I (Ang-I), and Apelin-13 peptide substrates (Figure 1b). We found wild-type ACE2-displaying yeast hydrolyze more than 95% of both Ang-II and Apelin-13 peptides during a 2.5-hour room temperature reaction, but do not yield measurable Ang-I hydrolysis. This outcome was expected given ACE2’s catalytic efficiency (k_cat_/K_m_) for Ang-I is nearly 1,000-fold lower than those for Ang-II and Apelin-13 [8]. We also tested our assay using a mixture of Ang-I, Ang-II and Apelin-13 peptides to understand ACE2’s behavior in a more realistic complex environment and to simultaneously profile activity toward multiple substrates. We found Ang-I acts as an inhibitor of ACE2-mediated Ang-II and Apelin-13 hydrolysis under these mixed substrate conditions, with Ang-II and Apelin-13 hydrolysis reduced approximately five-fold.

We next designed site-directed ACE2 libraries to target key residues controlling substrate preference. We examined the ACE2 crystal structure and a sequence alignment of related peptidases to identify ACE2 residues in the substrate binding pocket that display variation across homologs [16,17]. ACE2’s S_1_ subsite binds and recognizes the main portion of the peptide substrate N-terminal to the cleavage site, while the S_1_’ site accommodates the C-terminal residue (Figure 1c). We focused on residues R273, F504, Y510 and R514 from the S_1_ subsite and residues E145, N149, T347, M360, K363, and T371 from the S_1_’ subsite. We created ten individual degenerate NNB (N denotes all four nucleotides while B denotes C, G or T) codon libraries at each of these ACE2 positions.

We screened between five and fifteen clones for each of the ten site-directed ACE2 libraries using a mixed substrate assay (25 µM Ang-I, 25 µM Ang-II, 25 µM Apelin-13) to identify variants with improved Ang-II activity and specificity. A majority of the sampled clones from the R273, T347, F504, and R514 libraries had diminished Ang-II activity (Supplementary Table 1), while most of the clones from the E145, N149, and K363 libraries possessed substrate hydrolysis profiles similar to wildtype ACE2. The remaining M360, T371 and Y510 libraries all contained multiple clones with improved Ang-II hydrolysis and Ang-II:Apelin-13 specificity. We identified M360P, T371D, and Y510Ile as the top sampled single mutants at these three sites (Figure 1d).

We next constructed six second-generation libraries using M360P, T371D, and Y510Ile as parents: M360P with NNB at positions 371 or 510, T371D with NNB at 360 or 510, and Y510Ile with NNB at 360 or 371. We screened five randomly sampled clones from each library to assess the relative proportions of functionally impaired and improved variants (Supplementary Table 2). We found the M360P/T371NNB, Y510Ile/M360NNB and Y510Ile/T371NNB double mutant libraries warranted further screening by virtue of containing high fractions of active clones. We then screened 18-25 additional clones from these three libraries resulting in the four leading double mutants: M360P/T371S, M360P/Y510Ile, M360L/Y510Ile and T371L/Y510Ile (Figure 1d).

### Engineered ACE2s display enhanced kinetic properties

Our improved ACE2 variants were engineered in a yeast display format where they were fused to cell adhesion proteins on the yeast cell surface. We next wanted to fully characterize the ACE2 variants’ enzymatic activities in a soluble form similar to how they may ultimately be administered as therapeutics. We cloned wildtype ACE2 and the leading single and double mutants into a soluble secretion vector [18], expressed the enzymes in Human Embryonic Kidney (HEK) cells, and purified to enzymes to greater than 90% purity (Supplementary Figure 3). The enzyme purifications yielded approximately 1 mg/L HEK culture supernatant.

We performed initial enzyme activity assays and found any variant containing the M360P substitution had undetectable activity in the soluble form (Supplementary Figure 4). These results indicate our initial hits containing M360P were dependent on the yeast display fusion protein, and the activity increases did not translate to soluble ACE2 enzymes. We therefore did not further pursue any additional variants containing the M360P substitution.

We performed Michaelis-Menten kinetic studies on wildtype ACE2, Y510Ile, M360L/Y510Ile, and T371L/Y510Ile with the substrates Ang-II, Apelin-13, and Ang-I (Figure 2a). The three engineered variants displayed Ang-II turnover numbers (k_cat_) at least fourfold greater than wildtype ACE2, but also had increased Michaelis constants (K_m_). The T371L/Y510Ile variant had a 25% increase in Ang-II catalytic efficiency over wildtype. On the Apelin-13 substrate, the engineered variants showed wildtype-like turnover numbers, but also had 5-13-fold increased Michaelis constants, resulting in greatly diminished catalytic efficiencies. Wildtype ACE2 and the three variants all had very slow kinetics on the Ang-I substrate, indicating the new mutations did not introduce new activity on Ang-I.

**Figure 2.**
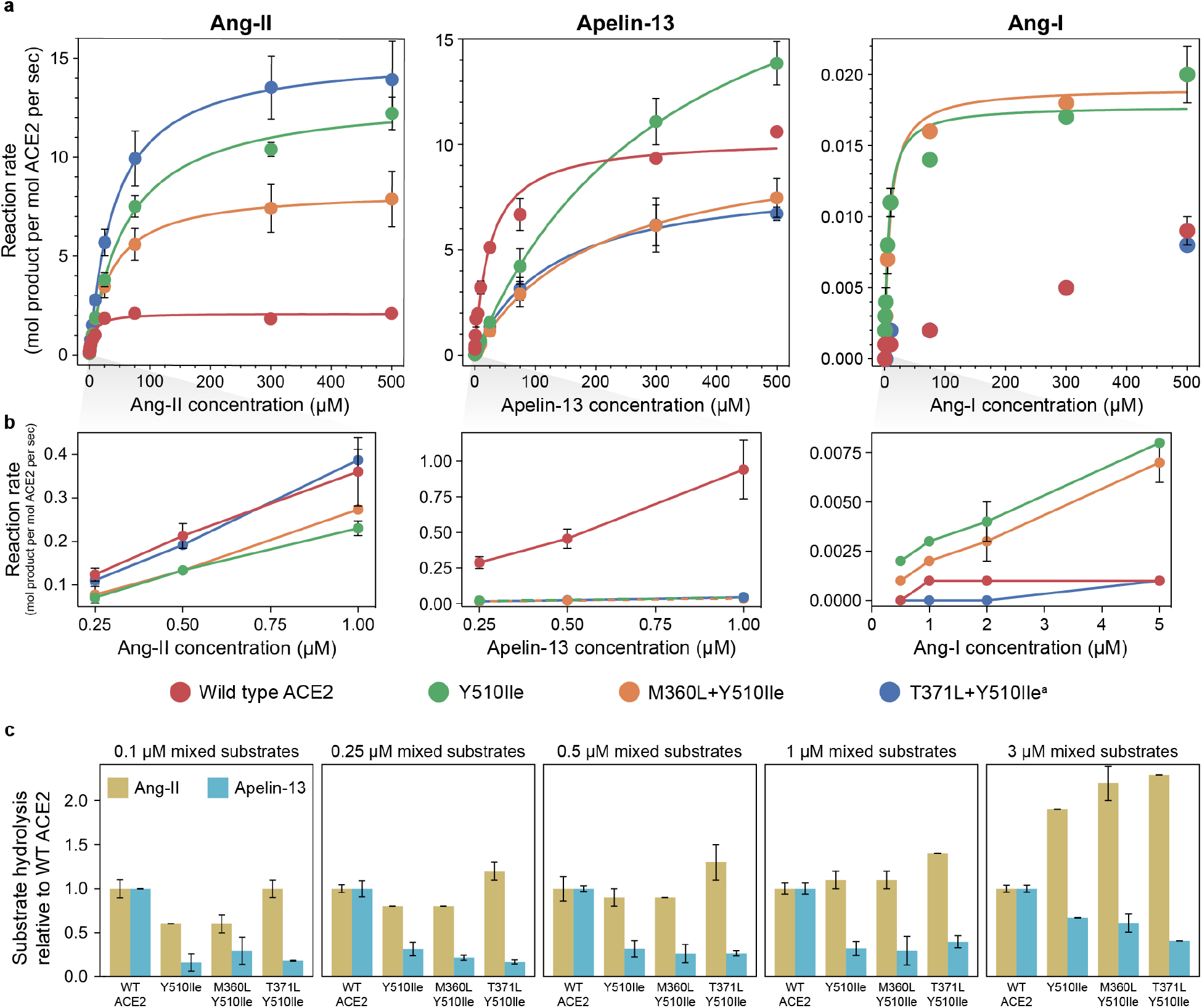
Kinetic properties of engineered ACE2 variants. (a) Initial rate plots for single substrate hydrolysis of Ang-II, Apelin-13, and Ang-I peptides. Error bars denote standard deviations for duplicate measurements and the absence of error bars for some data points indicates a standard deviation less than height of data point marker. The lines depict nonlinear regression fits to the Michaelis–Menten equation and the kinetic parameters are reported in Table 1. (b) Initial rate plots zoomed in to lower substrate concentrations that are relevant for physiological functioning *in vivo*. (c) Relative amounts of Ang-II and Apelin-13 hydrolysis products formed in mixed peptide substrate assays. Ang-I, Ang-II and Apelin-13 were present at identical concentrations in each reaction and incubated with 250 pM purified ACE2. Activity values are normalized to one for wild type ACE2. Error bars denote standard deviations (SD) for duplicate measurements. Error bars for wild type ACE2 values reflect SD in unnormalized hydrolysis product values for two trials. Error bars for mutant ACE2 values reflect SD in normalized hydrolysis product values. The slow rate of Ang-I hydrolysis relative to Ang-II and Apelin-13 hydrolysis precludes detection of the Ang-I hydrolysis product formation. ^a^ An additional P235Q mutation was observed in the T371L/Y510Ile variant due to polymerase error during cloning. This mutation is located at a surface position distal to the ACE2 active site and has been found to not affect enzyme activity or specificity.

We also evaluated the enzymes’ kinetic rates at very low substrate concentrations reflective of the physiological ranges found in human blood and tissues [11-13,19] (Figure 2b). The three engineered variants showed wildtype or near-wildtype-like Ang-II hydrolysis rates from 0.25 to 1 µM Ang-II. The engineered enzymes had practically no hydrolysis activity on Apelin-13 and were at least 10-fold lower activity than wildtype ACE2 from 0.25 to 1 µM Apelin-13. All reactions rates measured on Ang-I were 2-3 orders of magnitude slower than Ang-II hydrolysis, and are thus too slow to be of physiological relevance.

We evaluated wildtype ACE2’s and the engineered variants’ hydrolysis rates with equimolar mixtures of Ang-II, Apelin-13, and Ang-I to understand substrate preferences and potential interactions between substrates (Figure 2c). Across all mixed substrate concentrations tested, the leading ACE2 variant T371L/Y510Ile has Ang-II hydrolysis activity equal to or greater than wild type and a five-fold or greater decrease in Apelin-13 activity. Mixed substrate conditions can cause negative interactions between substrates when one substrate acts an inhibitor for another. We compared single and mixed substrate hydrolysis assays and found the presence of multiple substrates reduces hydrolysis rates for both Ang-II and Apelin-13 (Supplementary Figure 5). Ang-II hydrolysis rates are reduced between two-and three-fold while Apelin-13 hydrolysis rates are reduced by three to nine-fold. Given the results presented in Figure 1b, which indicate that Ang-I inhibits ACE2 hydrolysis of Ang-II and Apelin-13, it appears likely that the presence of Ang-I in the mixed substrate reactions is a substantial contributor to the observed reduction in Ang-II and Apelin-13 hydrolysis rates.

**Table 1.** Ang-II, Apelin-13 and Ang-I hydrolysis kinetic parameters for wild type and mutant ACE2s. Parameter estimates obtained by using nonlinear fits to initial rate data that appears in Figure 2a. N/D denotes not determined due to the maximum Ang-I concentration assayed (500 µM) being well below the concentration needed to reach the enzyme’s maximum hydrolysis rate; this phenomenon precludes accurate kinetic parameter estimation.

| ACE2 | Wild Type | Y510Ile | M360L/Y510Ile | T371L/Y510Ile |
| --- | --- | --- | --- | --- |
| Ang-II $k_{cat}$ ($s^{-1}$) | 2.1 +/- 0.2 | 13.2 +/- 0.7 | 8.4 +/- 0.1 | 15.3 +/- 0.3 |
| Ang-II $K_m$ (μM) | 7.2 +/- 2.8 | 60.4 +/- 10.7 | 35.9 +/- 2.1 | 42.1 +/- 2.5 |
| Ang-II $k_{cat}/K_m$ ($s^{-1}M^{-1}$) | $2.9 \times 10^5$ | $2.2 \times 10^5$ | $2.3 \times 10^5$ | $3.6 \times 10^5$ |
| Apelin-13 $k_{cat}$ ($s^{-1}$) | 10.3 +/- 1.2 | 23.1 +/- 0.6 | 10.4 +/- 0.3 | 8.6 +/- 0.3 |
| Apelin-13 $K_m$ (μM) | 25 +/- 10 | 330 +/- 20 | 200 +/- 12 | 130 +/- 15 |
| Apelin-13 $k_{cat}/K_m$ ($s^{-1}M^{-1}$) | $4.1 \times 10^5$ | $7.0 \times 10^4$ | $5.2 \times 10^4$ | $6.7 \times 10^4$ |
| Ang-I $k_{cat}$ ($s^{-1}$) | N/D | 0.018 +/- 0.02 | 0.019 +/- 0.01 | N/D |
| Ang-I $K_m$ (μM) | N/D | 6.4 +/- 2.9 | 8.6 +/- 2.5 | N/D |
| Ang-I $k_{cat}/K_m$ ($s^{-1}M^{-1}$) | N/D | $2.8 \times 10^3$ | $2.2 \times 10^3$ | N/D |

### Engineered ACE2s retain specificity against other off-target substrates

When administered as a therapeutic, ACE2 will encounter many potential substrates *in vivo* and we would like to minimize any off-target hydrolysis activity that may lead to undesirable physiological consequences. Our directed evolution experiments screened for variants with improved Ang-II activity and specificity over Apelin-13 and Ang-I. Without directly screening for all potential substrates, it’s possible our engineered variants could retain high activity on other peptide substrates recognized by ACE2 or even acquire activity on new substrates.

We tested our engineered ACE2 variants on additional peptides that were not assayed during the directed evolution process to further understand the substrate hydrolysis profiles. The Ang1-9 nonapeptide, DRVYIFPFH, is not hydrolyzed by wild type ACE2 [8]. We found none of the ACE2 variants exhibited activity toward this peptide as determined by LC analysis (Supplementary Figure 6). We also tested the variants’ activity on the known ACE2 substrate Des-Arg9-bradykinin (dR9-bk), an octapeptide with sequence RPPGFSPF [8,20] (Figure 3). All three engineered variants had strongly diminished activity on dR9-bk, despite not being directly evolved away from that substrate. Taken with the above observations of mutant ACE2 activities toward Ang-I and Ang1-9, these dR9-kb hydrolysis assay outcomes suggest that ACE2’s activity profile with respect to Ang-II and Apelin-13 hydrolysis can be enhanced without concomitant compromise of specificity toward off-target substrates other than Apelin-13 (Figure 3).

**Figure 3.**
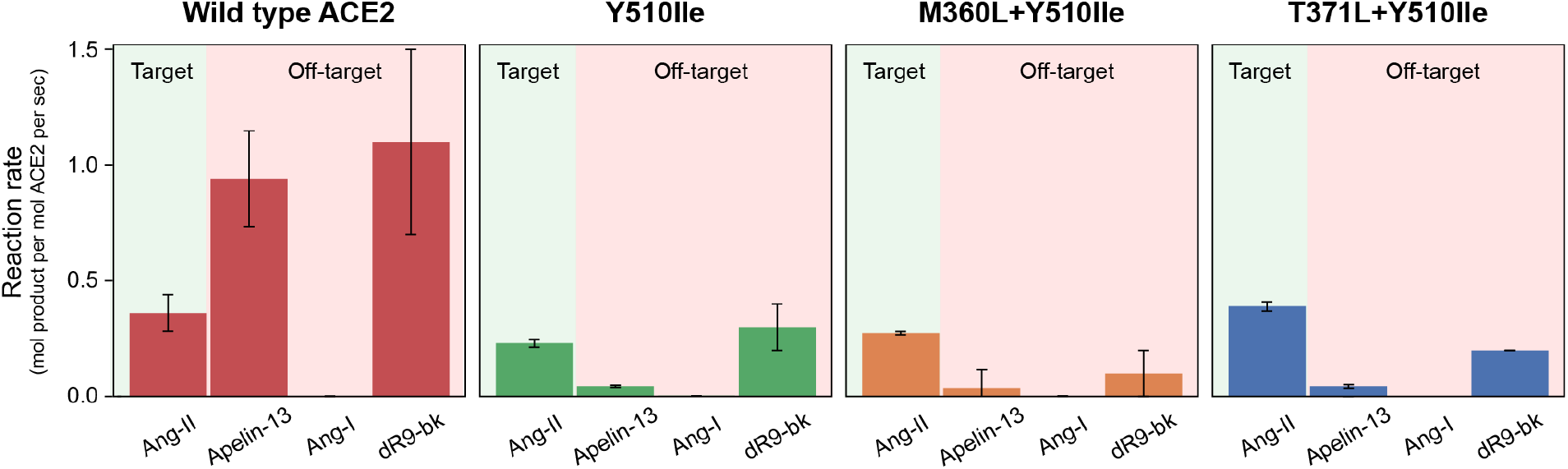
Peptide specificity profiles of engineered ACE2 variants. Purified ACE2s were incubated with 1 µM of peptide substrates in single substrate hydrolysis experiments. The engineered variants retain wildtype-like Ang-II hydrolysis, while having substantially reduced activity on Apelin-13 and dR9-bk. Error bars denote standard deviation for duplicate measurements.

### Structural rationale for enhanced activity and specificity

We examined the crystal structure of ACE2 [21] to postulate the mechanisms by which the M360L, T371L and Y510Ile substitutions influence ACE2’s hydrolytic activity and specificity. Y510 resides in the S1 subsite of ACE2’s substrate binding pocket (Supplementary Figure 7). ACE2 substrates, with the exception of the slowly-hydrolyzed Ang-I peptide, feature residues with small sidechains at the P1 position; both Ang-II and Apelin-13 carry proline at this position. The Y510Ile substitution may increase Ang-II hydrolysis by enlarging the S1 subsite and/or creating favorable hydrophobic interactions between ACE2 and the Ang-II proline sidechain. Ang-II and Apelin-13 have the same residues at the P1 and P1’ positions, and thus any specificity differences are likely to be influenced by interactions N-terminal to the P1 position. Y510Ile’s enhanced Ang-II specificity is either a result of direct interactions between Ile510 and these upstream substrate residues or possibly subtle conformational rearrangements that better accommodate the Ang-II substrate.

The sidechains of M360 and T371 line the interior of ACE2’s S1’ subsite. This subsite accommodates the C-terminal phenylalanine of both Ang-II and Apelin-13. The similarity in sidechain size among Met, Leu, and Thr suggests that sidechain hydrophobicity at these positions may contribute to the enhanced activity and specificity of M360L and T371L. Given that Ang-II and Apelin-13 are identical at the P_1_ and P_1_’ positions, there must be some indirect, longer-range effect, either caused by (1) differences in substrate positioning due to substrate residues N-terminal to P1 or (2) by structural changes induced by substitutions at sites 360/371.

## Discussion

ACE2 controls Ang-II levels in blood and tissues and this capability could be leveraged to treat a variety of human diseases. However, this enzyme is a central regulator of multiple peptide hormone signaling pathways and these varied physiological roles greatly limit its potential to specifically cleave Ang-II for therapeutic applications. In this work, we applied structure-guided directed evolution to engineer ACE2 variants to hydrolyze Ang-II with high activity and specificity.

Our leading ACE2 variant had the T371L and Y510Ile substitutions that resulted in a sevenfold increase in the Ang-II turnover number (k_cat_), a sixfold diminished catalytic efficiency (k_cat_/K_m_) on Apelin-13, and an overall decreased activity on other ACE2 substrates that were not directly assayed in the enzyme engineering work. The T371L/Y510Ile variant also has substantially increased Ang-II activity and specificity at low peptide substrate concentrations similar to those found in human blood and tissue. Importantly, the variant achieved these large functional changes with only two amino acid substitutions, making it nearly identical to wild-type ACE2 and reducing the likelihood that it would induce an undesired immune response when administered as a drug. All of the enzyme enhancements listed above suggest the T371L/Y510Ile variant would outperform wild-type ACE2 derivatives that have been and are currently being tested in clinical trials.

We developed a yeast surface display (YSD)-based workflow that enabled streamlined ACE2 variant expression, purification, and screening. This system allowed us to screen fifty ACE2 variants per day with detailed LCMS-based readouts of substrate hydrolysis profiles. One caveat with this approach is that it assays variants tethered to the yeast cell surface and therefore may discover variants that only function in this context. All variants containing the M360P mutation appeared as the top hits in the YSD screen but did not have detectable activity when expressed as soluble enzymes. This loss of hydrolytic activity observed for soluble ACE2 variants carrying the M360P substitution is not without precedent as display on the yeast surface has been observed to stabilize active conformations of protein mutants that are prone to taking on inactive conformations when free in solution [22,23]. This also illustrates the need to consider multiple diverse variants when validating the YSD screen with soluble enzyme expression assays. If we had simply chosen the top variants that contained the M360P mutation, the desirable effects of the T371L and Y510Ile substitutions in the context of soluble ACE2s would have been masked by the highly deleterious M360P mutation.

We performed our ACE2 variant screening with a mixture of three ACE2 substrates to simultaneously evaluate enzyme activity and specificity and to engineer ACE2 under conditions more similar to the *in vivo* environment. It also seems the mixed substrate screening provides a more stringent test of ACE2 activity because presence of multiple substrates reduces the individual reaction rates by several fold (Supplementary Figure 5). The mixed substrate conditions cause the ACE2 active site to be occupied by multiple different peptides that each effectively act as competitive inhibitors for the other substrates’ hydrolysis reactions. In addition to inhibition, the interactions between substrates also give rise to subtle shifts in substrate specificity due to the differing *k*_*cat*_ and *K*_*m*_ across different substrates. This highlights the importance of screening ACE2 variants under mixed substrate conditions that reflect the physiological environment.

This work performed an initial shallow screen to quickly identify ACE2 variants with improved activity and specificity. A more comprehensive screen of our ten current ACE2 single site saturation libraries and other site libraries would likely uncover ACE2 variants superior to our leading T371L/Y510Ile double mutant. The best mutant in each NNB site-directed library can be identified 95% of the time by screening only ninety clones [24]. Given our LC method’s screening capacity of approximately fifty clones per day, ninety clones from all ten active site saturation mutant libraries could be screened in less than one month. The leading single mutants could then be used to design focused double mutant libraries where one mutation is fixed and a second amino acid position is varied by introduction of a NNB codon. Twenty additional double mutant libraries could be screened with 95% coverage in less than two months. These deeper sequence space searches would almost certainly uncover variants that are markedly improved relative to T371L/Y510Ile.

The clinical potential of our engineered ACE2s could be further enhanced by increasing enzyme activity at physiologically relevant Ang-II concentrations, i.e, below 200 pM, and directly screening against known off-target ACE2 substrates including Apelin-13, des-Arg9-bradykinin, and Dynorphin-A. Follow up studies need to evaluate the pharmacokinetics to understand the retention time and distribution within mouse models, in addition to pharmacodynamics studies to identify the organs and tissues in which ACE2 is modulating the ATR biological axis. Carried out in parallel with toxicity studies, these animal experiments would open the door to *in vivo* evaluation of ACE2 variants in particular disease models as a prelude to human trials.

Our directed evolution method has enabled isolation of ACE2 variants with potential to outperform wild type ACE2 as protein therapeutics. This outcome is important for advancing current ACE2 therapeutic initiatives, such as treating respiratory distress associated with traumatic injuries or viral infection [4], and giving rise to new ACE2 biomedical applications including Alzheimer’s Disease therapy [25]. Furthermore, these results provide a foundation for developing additional directed evolution methods, such as microfluidics droplet-based screening to augment ACE2 activity and specificity [26], that facilitate sampling larger swaths of ACE2 sequence space and thus further enable ACE2 to realize its potential as an agent for treating and preventing a wide range of health conditions.

## Materials and Methods

### ACE2 library generation and screening

Residues 18-615 of the human (UniProt Q9BYF1) ACE2 genes were synthesized as a yeast codon-optimized gBlock (Integrated DNA Technologies, Coralville, IA) and ligated into the NheI and MluI sites of yeast display vector VLRB.2D-aga2 (provided by Dane Wittrup, MIT); this vector fuses the aga2 protein to the C-terminus of ACE2 (Supplementary Figure 1). Site-directed mutant libraries were constructed via overlap extension PCR and standard ligation using T4 DNA ligase (New England Biolabs, Beverly, MA). Oligonucleotide primers carrying NNB base triplets were purchased from Integrated DNA Technologies. The degenerate ‘B’ base is comprised of a mixture of C, G and T bases while the degenerate ‘N’ base is composed of all four nucleotide bases. Primers CDspLt (5’-GTCTTGTTGGCTATCTTCGCTG-3’) and CDspRt (5’-GTCGTTGACAAAGAGTACG-3’) were used as outer primers for ACE2 gene amplification reactions. Overlap extension PCR products were digested using NheI and MluI (New England Biolabs) and ligated into the yeast surface display vector VLRB.2D-aga2 digested with these same two enzymes. Overnight ligation reactions were desalted using a DNA Clean & Concentrate-5 column (Zymo Research, Orange, CA) and transformed into chemically competent NEB 5a cells (New England Biolabs).

For each transformation cells were plated onto three LB plates containing 100 mg/mL carbenicillin and after overnight incubation at 30°C all colonies on plates for each respective transformation were scraped into 10 mL of LB + carbenicillin liquid media and grown overnight at 30°C. Total colony counts for each single site-directed mutant library ranged from 500 to 2,000; numbers considerably greater than the fewer than 50 colonies observed for a negative control ligation reaction that contained digested VLRB.2D-aga2 backbone DNA absent any ACE2 gene insert.

DNA was harvested from overnight liquid cultures using the Qiagen Spin Miniprep Kit (Qiagen, Valencia, CA) and transformed into the EBY100 yeast surface display strain [14] that had been made chemically competent using the Frozen EZ-Yeast Transformation II Kit (Zymo Research). For each single site-directed mutant library transformed yeast were plated onto three each Synthetic Dropout minus Tryptophan (SD -Trp) plates and incubated at 30° C for two days. Individual yeast colonies were picked into 4mL of pH 4.5 Sabouraud Dextrose Casamino Acid media (SDCAA: Components per liter - 20 g dextrose, 6.7 grams yeast nitrogen base (VWR Scientific, Radnor, PA), 5 g Casamino Acids (VWR), 10.4 g sodium citrate, 7.4 g citric acid monohydrate) and grown overnight at 30°C and 250 rpm. For induction of ACE2 display a 5 mL pH 7.4 Sabouraud Galactose Casamino Acid (SGCAA: Components per liter - 8.6 g NaH_2_PO*H_2_O, 5.4 g Na_2_HPO_4_, 20 g galactose, 6.7 g yeast nitrogen base, 5 g Casamino Acids) culture was started at an optical density, as measured at 600 nm, of 0.5 and shaken overnight at 250 rpm and 20°C.

For LC determination of yeast-displayed ACE2 peptidase activity 35 µL of induced yeast culture was transferred to 500 µL of pH 7.4 Phosphate Buffered Saline (PBS) containing 0.1% (wt/vol) bovine serum albumin (BSA) and centrifuged at 4000g for one minute. Yeast were resuspended in 400 µL of ACE2 reaction buffer (50 mM 2-(N-morpholino)ethanesulfonic acid (MES), 300 mM NaCl, 10 µM ZnCl_2_, 0.02% w/v Hen Egg Lysozyme (Sigma-Aldrich, St. Louis, MO) as carrier protein, pH 6.5) containing 25 µM each Angiotensin-I (Anaspec, Fremont, CA), Angiotensin-II (Anaspec) and Apelin-13 (Bachem, Torrance, CA) and tumbled in microfuge tubes at 20 rpm at room temperature for 2.5 hours. Reactions were centrifuged at 9000g for one minute to pellet yeast and supernatants withdrawn for LC analysis.

Liquid chromatography (LC) analyses were performed on a Shimadzu 2020 LC-MS apparatus operating in UV detection mode at 214 nm. Sixty-five µL of ACE2 reaction were injected onto a 3×150 mm, 100 Å, 5 µm Polar C18 Luna Omega column (Phenomenex, Torrance, CA) using a linear gradient of 11% B to 100% B over 25 minutes with 5 minutes at final conditions and 8-minute re-equilibration. Mobile phase A consisted of 0.02% (v/v) trifluoroacetic acid in water, and mobile phase B consisted of 0.016% (v/v) trifluoroacetic acid in acetonitrile. Mobile phase flowrate was maintained at 0.3 mL/min throughout sample analysis. Hydrolysis product peaks were identified by comparison with chromatograms for reactions carried out using yeast displaying a negative control protein (horseradish peroxidase) and the area under the LC chromatogram curve, i.e, UV detector millivolts multiplied by elution timespan in seconds, was used as the metric for amount of product formed during the hydrolysis reaction.

For construction of ACE2 double mutant libraries that used leading single mutant clones, i.e., clone 360-7, 371-11, and 510-11 as described in the Results section, plasmids were rescued from 1 mL of yeast liquid culture using the Zymo Research Yeast Plasmid Miniprep II Kit. Rescued plasmids were used as templates for overlap PCR reactions carried out to introduce NNB codons, at the positions noted in the Results section, into the three respective single mutant ACE2 parent genes. Digestion and ligation of double mutant library PCR product DNA, *E. coli* transformation, *E. coli* culturing, double mutant library plasmid harvesting and EBY100 yeast transformations were carried out using procedures, as described above, identical to those for site-directed ACE2 single mutant libraries. Sequences for leading ACE2 double mutants were obtained by using the Zymo Yeast Plasmid Miniprep II kit, amplifying double mutant ACE2 genes using primers CDspLt and CDspRt, purifying the PCR products using a Zymo Clean & Concentrator-5 column and sequencing the PCR products using primers CDspLt, CDspRt and SqHmFw (5’-GGACTTTCAGGAAGACAACG-3’).

### Soluble expression & purification of wild type and ACE2 variants

Plasmid pcDNA3-sACE2(WT)-8his [18], which encodes human ACE2 residues 1-615 with native human codon representation and puts secretion of the gene product, with a C-terminal His8 tag, from mammalian cells under control of the cytomegalovirus CMV promoter, was used as template for creation of ACE2 genes carrying hydrolysis profile-modifying mutations identified during yeast displayed-ACE2 library screening. Mutant genes were constructed by overlap extension PCR using outer primers SqLtPck (5’-CGTGGATAGCGGTTTGACTCAC-3’) and PckRvSq (5’-CCTACTCAGACAATGCGATGC-3’), digested NheI-BamHI (New England Biolabs) and ligated into pcDNA3-sACE2(WT)-8his that had been digested using these same enzymes. Overnight ligation reactions were desalted, transformed into chemically competent NEB 5a cells as above and plated onto LB plates containing 100 mg/mL carbenicillin. Individual transformants were picked into 5 mL of LB media containing 100 mg/mL carbenicillin and incubated overnight at 37°C. DNA was harvested from overnight liquid cultures using the Qiagen Spin Miniprep Kit and inserted mutant ACE2 genes were verified by sequencing using primers SqLtPck, PckRvSq and PckF260 (5’-GTTACTGATGCAATGGTGGACC-3’).

Human Embryonic Kidney 293T cells (Product #CRL-3216, ATCC, Manassas, VA) were grown in DMEM (ATCC) supplemented with 10% Fetal Bovine Serum (FBS), 2mM L-glutamate (Invitrogen, Carlsbad, CA), and 1x Penicillin-Streptomycin (Invitrogen) in 100 mm tissue culture treated dishes under a 5% CO_2_ atmosphere at 37°C. Upon reaching ∼ 70% confluence cells were transfected with 7 µg of ACE2 expression plasmid DNA and 10 µL of jetPRIME transfection reagent (PolyPlus, Illkirch-Graffenstaden, France) per the manufacturer’s instructions. Two dishes of HEK cells were transfected for each ACE2 protein being expressed. Following the four-hour transfection interval DMEM containing DNA and transfection reagent was replaced with 10 mL of OptiMEM (Invitrogen) supplemented with 1x Pen-Strep and ACE2 expression was allowed to proceed for 40-48 hours at 37°C under 5% CO_2_.

Media was withdrawn from culture dishes, pooled for duplicate transfections, and centrifuged at 500g for 5 minutes. Supernatants were syringe filtered using 0.22 µm polyethersulfone (PES) Whatman PuraDisc filters (Cytiva, Marlborough, MA). Supernatant concentration and buffer exchange into pH 7.4 PBS was performed using 30-kDa MWCO VivaSpin20 centrifugal concentrator units (Cytiva). His_8_-tagged ACE2 proteins were purified from concentrated, buffer exchanged supernatants. Concentrated supernatants were brought to a final volume of 1 mL by addition of pH 7.4 PBS, adjusted to an imidazole concentration of 10 mM and tumbled head-over-head at 20 rpm in microfuge tubes with 25 mL of Qiagen Ni-NTA resin for 1 h at room temperature. Resin was subsequently washed with 1 mL PBS and 1 mL PBS/15 mM imidazole prior to ACE2 elution via 15-minute room temperature incubation of resin in 140 mL of pH7.4 PBS containing 250 mM imidazole. Eluted ACE2 proteins were buffer exchanged into 25mM Tris-HCl, pH 7.5 containing 10 µM ZnCl_2_ using Zeba Spin desalting columns (Fisher Scientific, Waltham, MA) and protein concentrations determined using the Pierce Coomassie Protein Assay kit (Fisher Scientific).

SDS-PAGE analysis of purified ACE2s was carried out using a LIfeTech Mini Gel Tank electrophoresis system. Approximately 1 mg of purified, desalted ACE2 was loaded into each well of a Novex 4-20% Tris-Glycine gel and electrophoresis was performed at 200 volts for 45 minutes. PageRuler Unstained Protein Ladder (Fisher Scientific) was used as a molecular weight standard. Gels were stained with Simply Blue SafeStain (Invitrogen) following electrophoresis.

### Activity and specificity profiling of purified ACE2 proteins

Peptide substrate hydrolysis assays were carried out at room temperature and 150 µL scale in 96-well plates using 50 mM MES, 300 mM NaCl, 10 µM ZnCl_2_, 0.01% (vol/vol) Brij-35 (Sigma-Aldrich) as reaction buffer. Reactions were halted by addition of 15 µL of 1M EDTA solution, pH 8, after intervals, which ranged from three minutes to 2.5 hours, during which less than 15% of the input peptide substrate had been hydrolyzed. ACE2 concentrations added to assays for the various peptide substrates evaluated were as follows: Ang-I/1.75 nM, Ang-II/700 pM, Apelin-13/1.4 nM, des-Arg9-bradykinin/700 pM, Ang (1-9)/700 pM, Multiplex/250 pM. Des-Arg9-bradykinin and Ang1-9 were purchased from Anaspec and ApexBio (Houston, TX) respectively.

Moles of product formed in hydrolysis reactions used for ACE2 activity and specificity profiling were determined by comparing area under the curve for LC chromatograms corresponding to these reactions to areas under the curve observed for control reactions in which > 95% input substrate conversion was achieved by incubation of substrate with 25 nM recombinant human ACE2 commercial standard (BioLegend, San Diego, CA) for between one and four hours pending peptide substrate. Elution times for peptide substrate inputs and hydrolysis products were determined by comparing chromatograms for the 25 nM ACE2 commercial standard reactions to input peptide reaction mixtures to which no ACE2 was added. LC analysis injection volumes and mobile phase gradient parameters were identical to those described above for yeast-displayed ACE2 library screening. Kinetic parameters for Ang-I, Ang-II and Apelin-13 hydrolysis were determined by fitting initial rates data using GraphPad Prism software (GraphPad, San Diego, CA).

## Acknowledgements

This work was supported by the US National Institutes of Health (5R35GM119854). “The content is solely the responsibility of the authors and does not necessarily represent the official views of the National Institutes of Health.”

