## Supplement for "Directed evolution of angiotensin-converting enzyme 2 (ACE2) peptidase activity profiles for therapeutic applications"

**Supplementary Information**

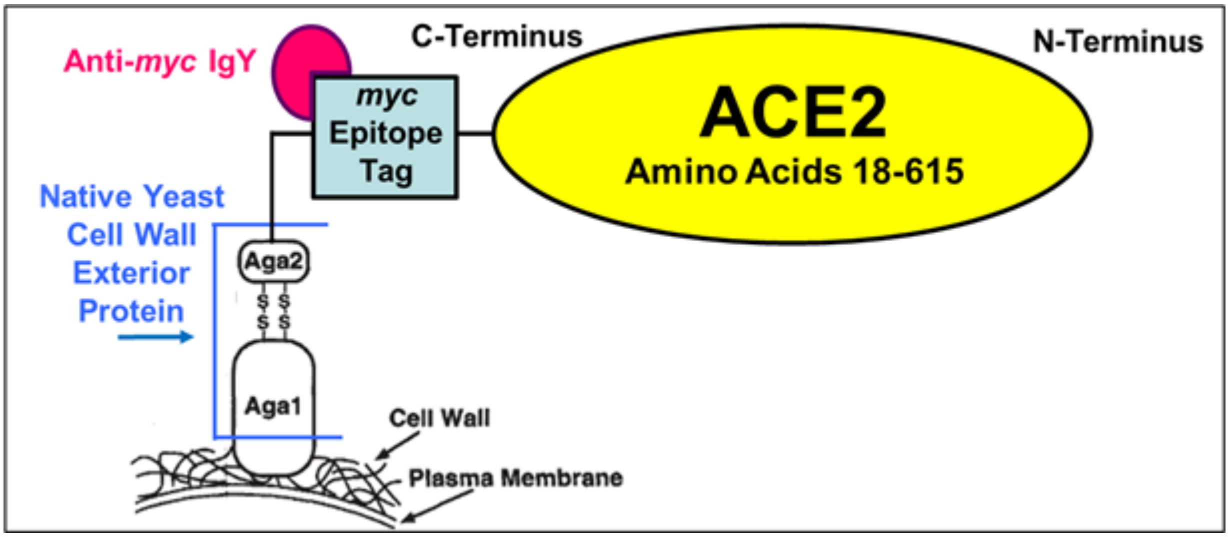


**Supplementary Figure 1.** Yeast display schematic. We fused ACE2's C-terminus to the Aga2 native yeast surface protein on yeast cell wall. We quantified ACE2 display using an anti-*myc* IgY (from chicken) in combination with an Alexa488-conjugated anti-chicken IgG (from goat) as secondary label . Each yeast cell displays up to $10^4$ copies of a single ACE2 variant on its surface. Schematic adapted from Boder and Wittrup, *Nat Biotechnol.* 1997 15(6):553-7.

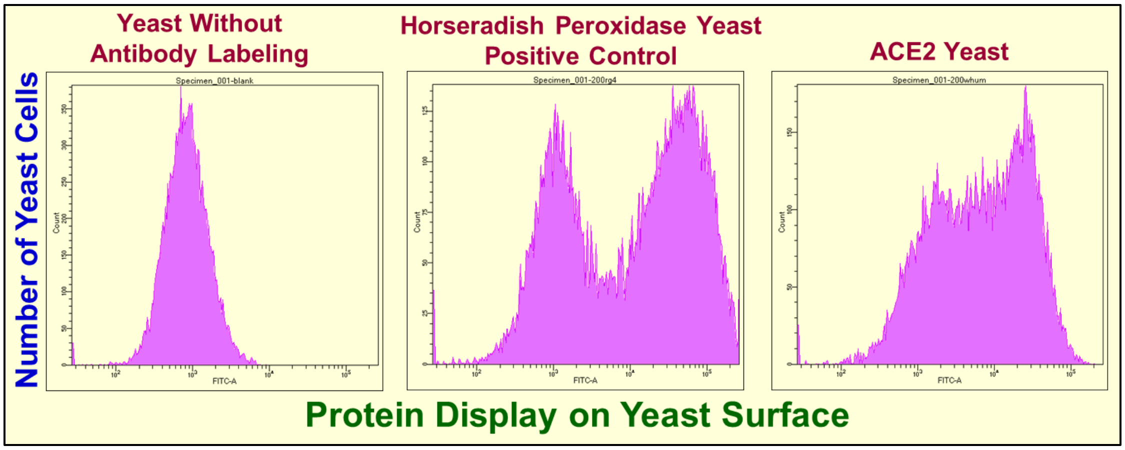


**Supplementary Figure 2.** Histograms for flow cytometric analysis of ACE2 display on yeast surface. One-hundred thousand yeast cells were incubated for one hour at 4°C on a tube rotator at 18 rpm in 200 µL of PBS/0.2% BSA containing 3 µg/mL anti-*myc* IgY (Aves Labs, Tigard, OR). Following primary labeling yeast were washed once in PBS/0.2% BSA and rotated at 18 rpm in same buffer containing 2 µg/mL Alexa488-conjugated goat anti-chicken IgG (Jackson ImmunoResearch, West Grove, PA). Histogram X-axes denote Alexa488 fluorescence (ACE2 display); Y-axes denote number of yeast cells with a given level of fluorescence. For poorly understood biological reasons homogeneous populations of yeast carrying identical display plasmids feature 25% or greater cells (left of histograms) that do not display protein. Positive control yeast carry display plasmid for horseradish peroxidase. Histograms depict results for $1.5*10^4$ yeast cells. Data collected using a Becton-Dickinson Fortessa X-20 flow cytometer.

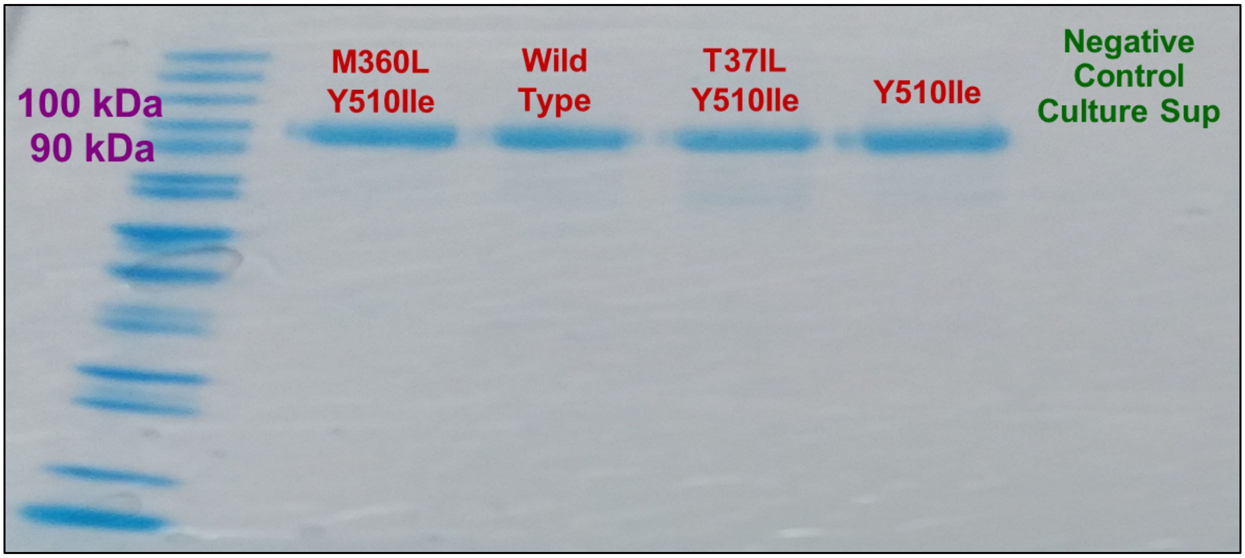


**Supplementary Figure 3.** Post-purification SDS-PAGE analysis of wild type and leading mutant ACE2s. Empty vector transfection culture negative control supernatant appears in rightmost lane. Expression levels and purities for the M360P and M360P/Y510 Ile mutants, which had low Ang-II hydrolysis activities and are not included in this gel, were similar to those for wild type ACE2.

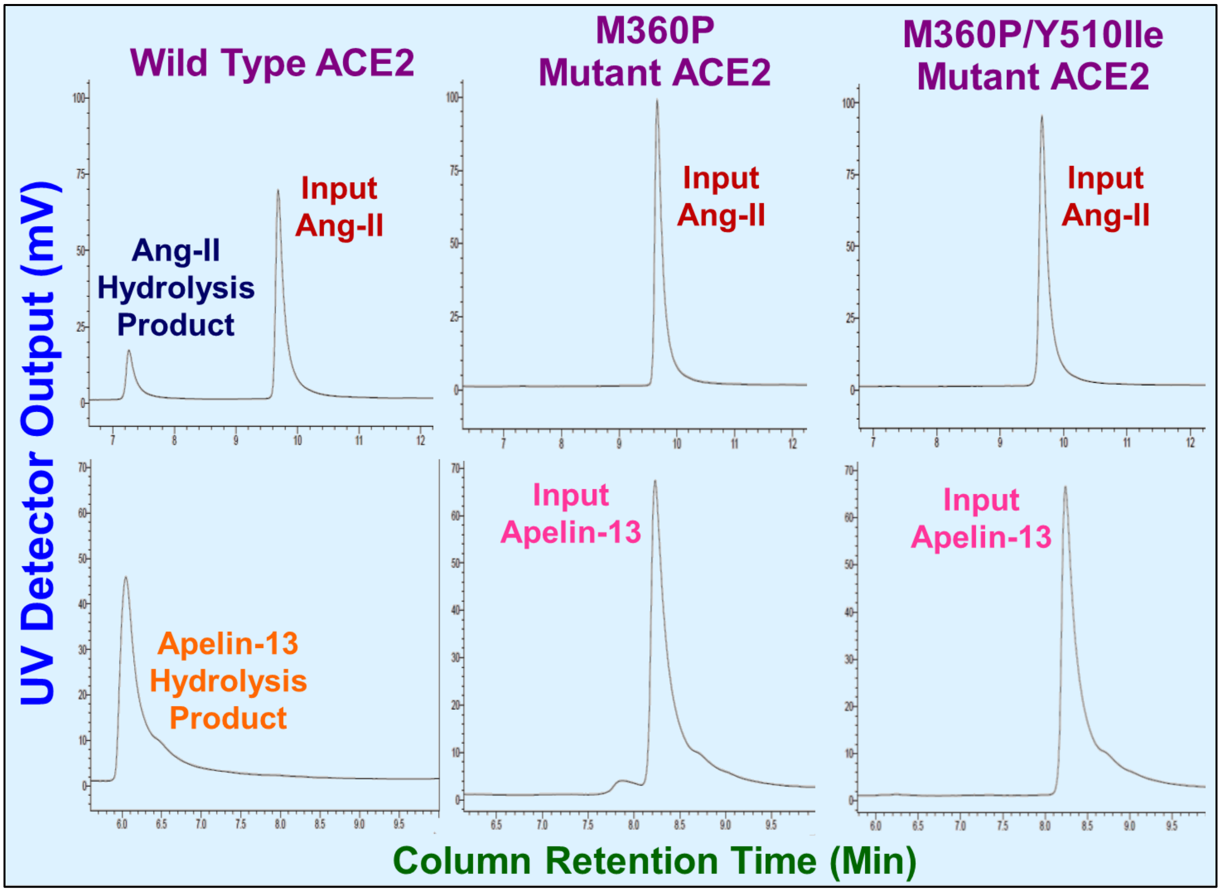


**Supplementary Figure 4.** LC chromatograms showing absence of hydrolytic activity for M360P and M360P/Y510Ile ACE2 mutants when expressed as soluble enzymes. Upper panel: Single substrate hydrolysis assay with Ang-II peptide (25 μM peptide, 2 nM purified ACE2). Input Ang-II peptide peak at ~ 9.8 min, hydrolysis product peak at ~ 7.3 min. Lower panel: Single substrate hydrolysis assay with Apelin-13 peptide (25 μM peptide, 15 nM purified ACE2). Input Apelin-13 peptide peak at ~ 8.3 min, hydrolysis product peak at ~ 6.1 min. Hydrolysis reactions were run for 2.5 hrs at room temperature.

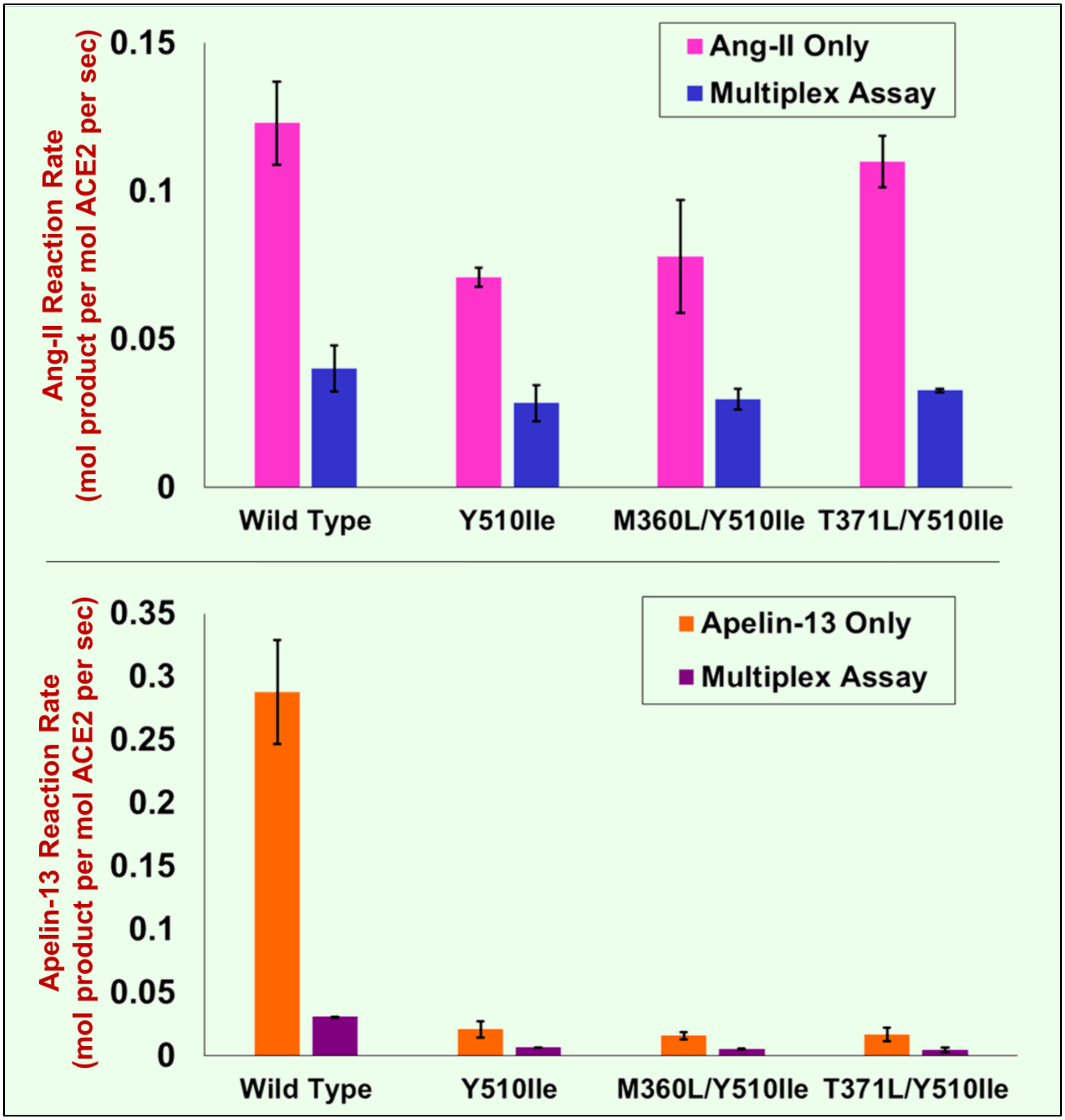


**Supplementary Figure 5.** Comparison of mutant and wild type ACE2 Ang-II (upper panel) and Apelin-13 (lower panel) hydrolysis rates across single substrate and mixed substrate assays carried out with peptide concentrations of 0.25 μM. Error bars denote standard deviations for duplicate measurements.

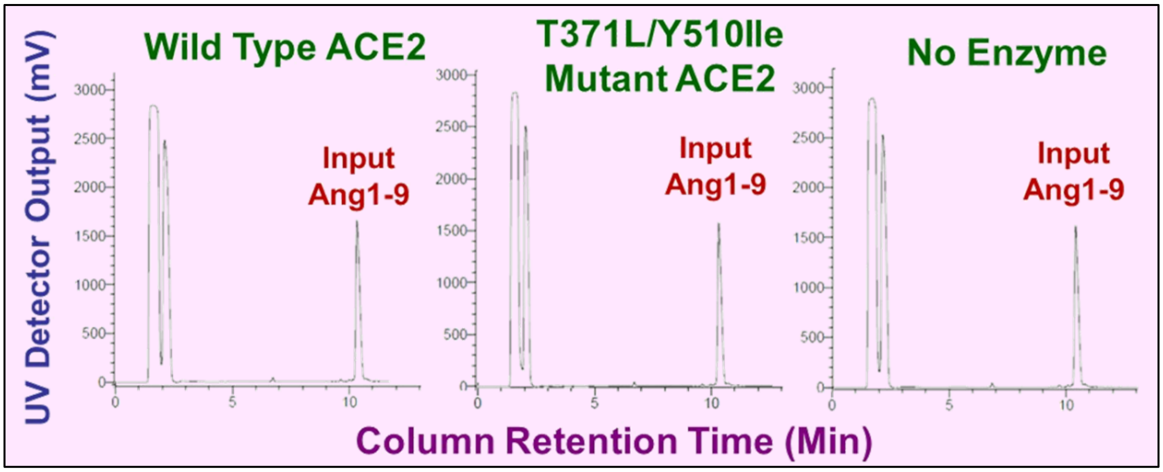


**Supplementary Figure 6.** LC chromatograms for Ang1-9 hydrolysis assays. Analyses correspond to assays carried out with 100 *u*M Ang1-9 peptide. Peaks appear for the input Ang1-9 peptide at ~ 10.2 min). No visible peak for Ang1-9 hydrolysis product; C-terminal hydrolysis of Ang1-9 would yield Ang-II and be expected to elute from the LC column at approximately 8.8 minutes. Peaks in 2-3 minute range are $Zn^{2+}$-EDTA complexes that form after EDTA is added to chelate the ACE2 active site $Zn^{2+}$ ion and thus halt any ongoing substrate hydrolysis.

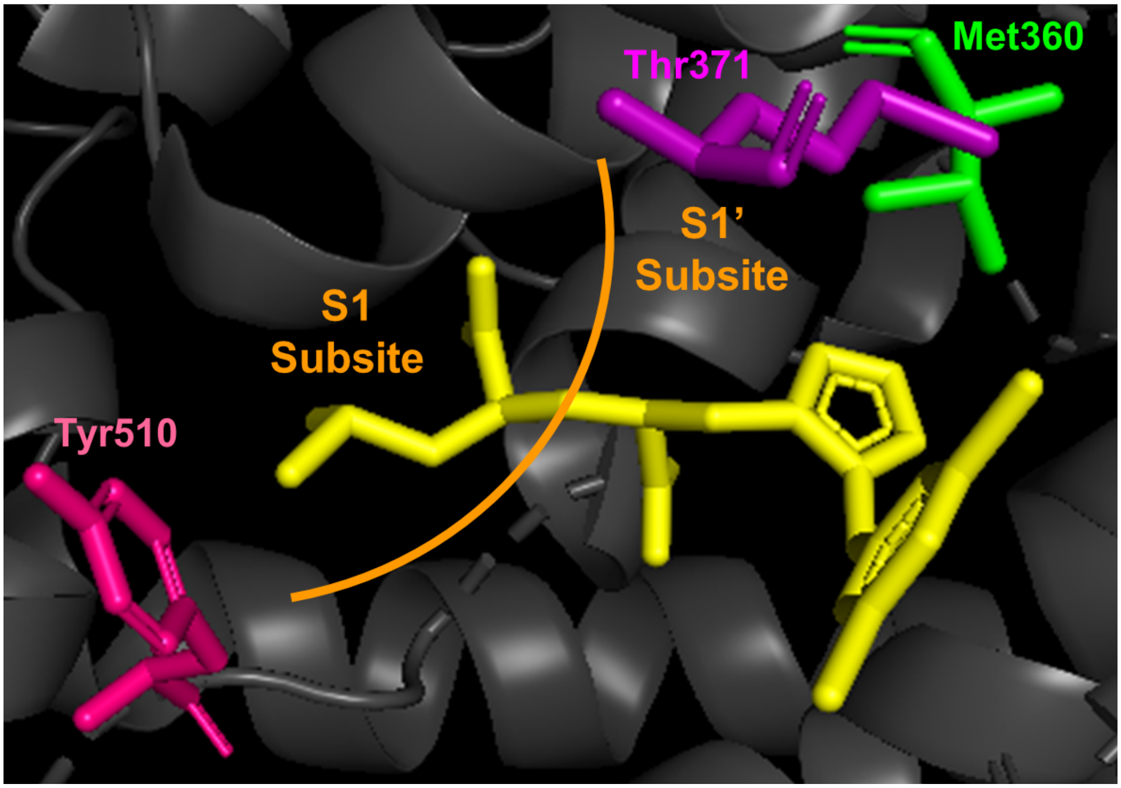


**Supplementary Figure 7.** Positioning of key residues Met360 (green), Thr371 (purple) and Tyr510 (magenta) in ACE2 substrate binding pocket (pdb entry 1R4L). Co-crystallized ACE2 small molecule inhibitor denoted in yellow. Orange arc denotes approximate boundary between upstream S1 subsite and downstream S1' subsite. Selected ACE2 residues and catalytic $Zn^{2+}$ ion hidden from image to provide enhanced view of M360, T371 and Y510 ACE2 residues and small molecule inhibitor. Figure created using Pymol.

**Supplementary Table 1.** Peptide hydrolysis profiles for clones characterized during initial LC screening of ACE2 site-directed single mutant libraries. Functionally impaired clones defined as ACE2 clones with less than 50% wild type ACE2 activity toward both Ang-II and Apelin-13 peptides. Improved clones defined as ACE2 variants with greater than 50% wild type ACE2 activity toward Ang-II and Ang-II:Apelin-13 hydrolysis ratio increased two-fold relative to the ratio for wild type ACE2.

| ACE2 Position Mutated | % Functionally Impaired Clones | % Clones Similar to Wild Type | % Improved Clones |
|---|---|---|---|
| E145 | 0/5 (0%) | **5/5 (100%)** | 0/5 (0%) |
| N149 | 0/5 (0%) | **5/5 (100%)** | 0/5 (0%) |
| R273 | **4/5 (80%)** | 1/5 (20%) | 0/5 (0%) |
| T347 | **3/5 (60%)** | 2/5 (40%) | 0/5 (0%) |
| M360 | 2/12 (17%) | 6/12 (50%) | **4/12 (33%)** |
| K363 | 0/5 (0%) | **5/5 (100%)** | 0/5 (0%) |
| T371 | 0/2 (0%) | 10/12 (83%) | **2/12 (17%)** |
| F504 | **5/5 (100%)** | 0/5 (0%) | 0/5 (0%) |
| Y510 | 7/15 (47%) | 0/15 (0%) | **8/15 (53%)** |
| R514 | **7/10 (70%)** | 3/10 (30%) | 0/10 (0%) |

**Supplementary Table 2.** Peptide hydrolysis profiles for clones characterized during initial LC screening of ACE2 site-directed double mutant libraries. Functionally impaired clones defined as ACE2 clones with less than 50% wild type ACE2 activity toward both Ang-II and Apelin-13 peptides. Improved clones defined as ACE2 double mutants with greater than 50% parental single mutant activity toward Ang-II and Ang-II:Apelin-13 hydrolysis ratio increased two-fold relative to the ratio for the parental single mutant.

| ACE2 Double Mutant Library | % Functionally Impaired Clones | % Clones Similar to Single Mutant Parent | % Improved Clones |
|---|---|---|---|
| M360P/NNB 371 | 2/5 (40%) | **3/5 (60%)** | 0/5 (0%) |
| M360P/NNB 510 | **4/5 (80%)** | 1/5 (20%) | 0/5 (0%) |
| T371D/NNB 360 | **5/5 (100%)** | 0/5 (0%) | 0/5 (0%) |
| T371D/NNB 510 | **5/5 (100%)** | 0/5 (0%) | 0/5 (0%) |
| Y510Ile/NNB 360 | 2/5 (40%) | **3/5 (60%)** | 0/5 (0%) |
| Y510Ile/NNB 371 | 0/5 (0%) | **4/5 (80%)** | **1/5 (20%)** |
